# Sex-specific effects of chronic paternal stress on offspring development are partially mediated via mothers

**DOI:** 10.1101/2022.03.25.485798

**Authors:** Rahia Mashoodh, Ireneusz B. Habrylo, Kathryn Gudsnuk, Frances A. Champagne

## Abstract

Paternal stress exposure is known to impact the development of stress-related behaviors in offspring. Previous work has highlighted the importance of sperm mediated factors, such as RNAs, in transmitting the effects of parental stress. However, a key unanswered question is whether mothers’ behavior could drive or modulate the transmission of paternal stress effects on offspring development. Here we investigate how chronic variable stress in Balb/C mice influences the sex-specific development of anxiety- and depression-like neural and behavioral development in offspring. Moreover, we examined how stressed fathers influenced mate maternal investment towards their offspring and how this may modulate the transmission of paternal stress effects on offspring. We show that paternal stress leads to sex-specific effects on offspring behavior. Males that are chronically stressed sire female offspring that show increased anxiety and depression-like behaviors. However, male offspring of stressed fathers show reductions in anxiety- and depression-behaviors and are generally more exploratory. Moreover, we show that females mated with stressed males gain less weight during pregnancy and provide less care towards their offspring which additionally influenced offspring development. These data indicate that paternal stress can influence offspring development directly and indirectly via changes in mothers, with implications for divergent development between male and female offspring.

## Introduction

It is well-acknowledged that the life-histories and experiences of parents prior to conception have a significant influence on behavioral and physiological development with effects that can persist for a number of generations (1,2). This phenomenon has been reported to occur in response to a broad range of physiological and social challenges, including but not limited to, drug/chemical exposures (3), dietary/immune challenges (4–6) and psychological stressors (7–9). For example, studies in laboratory rodents indicate that both maternal and paternal stress lead to alterations in the hormonal and behavioral response to stress of offspring (8,10–12). These findings highlight the importance of environmental factors in conferring disease risk and/or resilience. While the phenomenon of these parental effects is well supported, the mechanisms that account for these effects is a topic of speculation within the scientific literature and likely involves a complex interplay of molecular, physiological and behavioral factors.

In mammals, the intergenerational influence of mothers is likely to involve pre- and postnatal maternal interactions that have developmental programming effects on offspring. However, the phenomenon of paternal effects, particularly in species in which there is limited or no post-conception interaction between fathers and offspring is suggestive of a germline mechanism of inheritance. Indeed, there has been a significant focus on sperm-mediated mechanisms (*e.g.,* sperm RNAs and DNA methylation) in driving the effects of paternal stress (8,11–14). Our previous work, using embryo transfer, highlights the role of sperm-mediated factors and suggests that paternal pre-conceptual stress can predict offspring neurobiological and behavioral phenotypes. However, this work also highlights how pre- and postnatal maternal interactions may influence these outcomes in addition to any epigenetic effects in the sperm (4).

A role for maternal effects in mediating or moderating the impact of fathers on offspring is not typically considered within molecular studies of paternal transgenerational effects. However, the concept that females may dynamically adjust the provision of pre- and postnatal care towards offspring based on the quality of male mates has been well-acknowledged in the behavioral ecology literature and described across several non-mammalian species [*e.g.,* birds, insects, fish *etc.* (15–20)]. For example, females can increase investment in offspring sired by attractive and/or high-quality mates [or decrease investment in less desirable mates; termed differential allocation; (15,20)]. Alternatively, females could compensate by increasing investment in offspring of low quality mates [termed reproductive compensation;(16–18)]. It remains unclear what factors predict whether differential allocation or reproductive compensation will occur, though it likely involves a combination of the features of the male phenotype and female reproductive and energetic states (18,20,21). In mice, these maternal contributions could come in the form of prenatal investment (*e.g.,* increased feeding), maternal care (licking/grooming and nursing) and even from mothers’ microbiomes (4,22,23).

We have previously shown that differential allocation can occur in Balb/C mice mated with socially enriched males (23). Further, we have demonstrated that female mice mated with food restricted males show increased maternal investment in offspring (e.g. prenatal weight gain, postnatal maternal behavior) which buffered against the negative consequences predicted by paternal food restriction (4). Thus, paternal effects could be exacerbated or buffered depending on how mothers are impacted by the phenotype of fathers. Despite an expanding literature on the non-genetic transmission of paternal effects, we still do not have a clear understanding of the extent to which paternally-induced maternal effects occur across qualitatively different pre-conceptual exposures. In the current study, we further explore this phenomenon by examining the impact of variable and chronic pre-conceptual stress of adult Balb/C male mice to parse out direct (paternal) and indirect (maternal) effects on the subsequent development of anxiety- and depression-like behavior in offspring.

## Methods

### Animals, Husbandry & Breeding

Adult male and female Balb/C mice (F0; approximately 3 months of age) purchased from Charles River were used to generate offspring for these studies. Mice were housed on a 12-hour dark-light cycle at the Department of Psychology at Columbia University, with lights on at 22:00 and off at 10:00. All animals were given ad libitum access to food (mouse chow) and water. Adult male and female mice were housed in same-sex quads in 35 x 21 x 14cm Plexiglas cages in the animal facility for 2 weeks prior to mating. All procedures were conducted in accordance with animal care standards and approval of the Columbia University Institutional Animal Care and Use Committee (IACUC).

### Paternal Chronic Stress Paradigm

Adult male mice (N=12/group) were chronically stressed (stressed; PS) for a 6-week period during which each mouse was either exposed daily to a 1-h restraint stressor or a 6-min forced swim. The timing of the stressor was varied each day with a rest day interspersed every 4-5 days. Male mice were tested for anxiety- and depression-like behavior in the open-field and forced swim tests, respectively, exactly one week after the last stress exposure. Control mice were left undisturbed except for weekly cage changes (control; PC).

### Mating

A single male was placed in a mating group with 3 adult (6-8 week old) Balb/C female mice for approximately 2 weeks. After the mating period, males were removed and once females reached late pregnancy they were separated and singly housed prior to parturition (N=72). Changes in pre- and postnatal maternal investment were measured across gestation and during the first postnatal week for all litters (described below). At birth, pups were weighed and counted but otherwise left undisturbed during the postnatal period. All litters were observed from PN1-6 to determine postnatal levels of maternal care (frequency of licking/grooming, nursing). Following the final maternal observation on PN6, litters were weighed and counted but otherwise left undisturbed with the exception of weekly cage cleaning until weaning (PN28). At weaning, individual pups were weighed and placed into same-sex groups of four. From each litter, a maximum of 2 male and female offspring were selected for behavioral testing for a total of N=15/group/sex.

#### Prenatal Maternal Investment

As a proxy measure for prenatal investment (*e.g.,* food consumption during gestation), female mice that mated with PS or PC males were weighed daily across gestation (as previously described in (4). The day of birth was considered postnatal day 0 (PN0) and therefore, assuming an average gestation time of 19 days, percent weight gain was calculated for the last 20 weight observations. Given that there is individual variation in body weight, percent weight gain for each gestational day (*gd*) was calculated by subtracting current weight from initial weight (*w*_0_) and dividing by initial weight and multiplied by 100.

#### Postnatal Maternal Investment

Following parturition, dams were observed to determine whether mating condition results in variation in postnatal maternal behaviors. The procedure for assessing maternal behavior in mice has been described previously (24). Each dam was observed for four 1h periods per day by an observer blind to paternal condition from PN1-6, resulting in a total of 480 observations of each litter. The frequency of the following behaviors was scored: mother in contact with pups, mother in nursing posture over pups and mother licking and grooming any pups (N=21 and N=34 for PS and PC mated, respectively).

### Behavioral Testing of Offspring

Males exposed to stress or control conditions and male and female offspring (starting at PN55) from the four groups (N=15/group/sex) underwent testing in the open-field and forced swim test.

#### Open Field Test

The open field apparatus used was a 60 x 60 x 40cm Plexiglas box with black walls and a white floor. On the day of testing, the mouse was removed from its home cage and placed directly into one corner of the open field. After a 10-min session, the mouse was returned to its home cage. All testing was conducted under red lighting conditions. Behavior in the apparatus was video recorded. Behaviors scored using Ethovision (Noldus) included: (1) center area exploration, defined as the time spent in the inner (30 x 30cm) area, (2) latency to enter the center area, and (3) total distance travelled.

#### Forced-Swim Test

Depression-like behavior was measured during a brief forced-swim test. All forced-swim tests were conducted during the dark cycle in white light illuminated room. Mice were placed into a 2L glass beaker filled with water at room temperature (approximately 25 ± 2°C). All tests were video recorded for later scoring by an observer blind to condition. The behaviors scored were active struggling (vigorous swimming), and immobility (passive swimming, little to no active movement).

### Quantitative Real-Time PCR Analysis

RNA was isolated from the PN6 and adult hypothalamus of male and female PS and PC offspring using the AllPrep DNA/RNA Mini Kit (Qiagen) and reverse transcribed to cDNA using the SuperScript III First-Strand Synthesis System for RT-PCR applications (Invitrogen). Quantitative RT-PCR was performed with 1μl of cDNA using an ABI 7500 Fast Thermal Cycler and the Fast SYBR Green Master Mix reagent (Applied Biosystems). Primer probes (Sigma-Aldrich) were designed to span exon boundaries ensuring amplification of only mRNA (see **Table S1**). For each gene, C_T_ values were normalized to cyclophillin A (endogenous control). Relative expression values were obtained by the DDC_T_ method calculated relative to control group.

### Statistical Analyses

All statistical methods were performed using custom scripts written in *R* (version 3.5.1;(25)). Data wrangling and visualization was performed using a combination of base functions and the ‘tidyverse’ suite of R packages (26). Analysis of male behavior in response to chronic stress was performed using the base stats package in R. To account for the multilevel structure of the data (*i.e.*, male mice mated with multiple females and sired multiple litters and multiple offspring from the same father) linear multilevel mixed regression models were used where appropriate with either male and/or female ID included in the model as a random effect. These analyses were performed using the lme4 and lmerTest R packages (27,28). Bootstrapped mediation analyses were performed using the ‘mediation’ package in R (29).

## Results

### Effects of chronic stress on male behavior

#### Open-Field Test

Stressed adult male mice showed increased anxiety-like behavior when tested in an open-field. Stressed male mice (PS-F0) had a longer latency to enter the center (F(1,22)=7.999, p=0.00979), made less frequent entries into the center (F(1,22)=6.54, p=0.018) and spent less overall time in the center of the arena (F(1,22)=4.378, p=0.048; **Figure 1a**). There were no significant differences in number of fecal boli deposits (F(1, 22) = 1.042, p=0.318) or in the total distance travelled (F(1, 22)=1.678, p=0.209) between the two groups.

**Figure 1.**
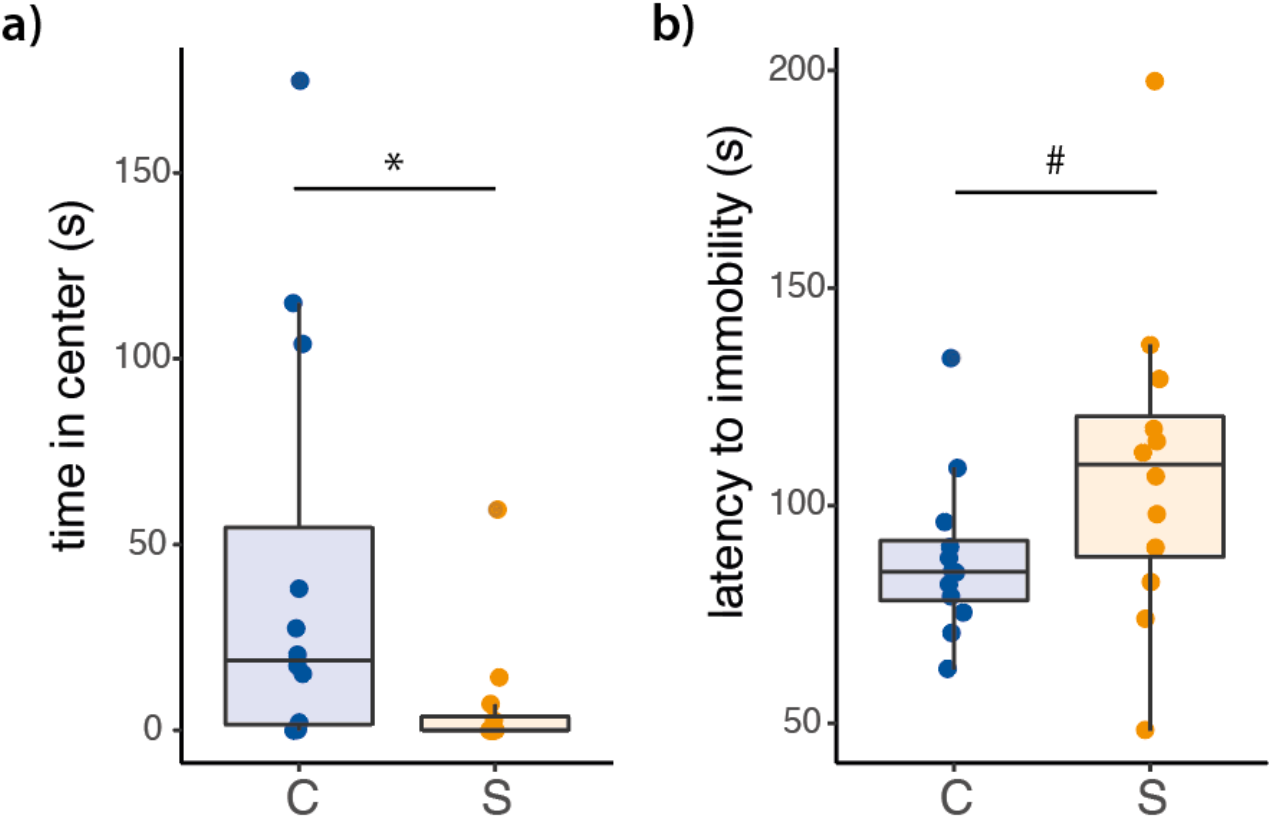
Adult male mice exposed to chronic variable stress (S) **(a)** spend less time in the center of an open-field test and **(b)** more time immobile in a forced-swim test compared with control (C) males (*\*p<0.05; #p<0.1*).

#### Forced-Swim Test

There was a trend for stressed males (PC-F0) to have an increased latency to immobility (t(21)=1.742, *p*=0.09; **Figure 1b**). Otherwise, there were no other major significant differences in forced-swim behavior between PS-F0 and PC-F0 males, including duration of time spent immobile during the last 4m of the test (t(21)= −1.397, p=0.1770).

### Maternal investment of females mated with stressed males

There was a trend for females that mated with stressed males to be less likely to become pregnant or maintain a successful pregnancy with only 36% of PS-mated females giving birth compared to 62% of PC-mated females (Survival Analysis, *p=*0.09; **Figure 2a**). There were no effects of paternal stress on litter size (Beta=−0.3967, t(37)=−0.420, p=0.677). There was a trend for stressed fathers to sire smaller litters (after controlling for litter size; beta=−0.475, t(36)=−1.719, *p=*0.09). Among females that became pregnant and successfully gave birth, we found that females mated with a PS male gained less weight in the final days of gestation. Among females that became pregnant and gave birth, there were significant differences in weight gain across the last 3 days of gestation between females mated with PS-F0 and PC-F0 males (beta = −5.1413, t(94)=−2.136, *p*=0.0353; **Figure 2b**). These effects were present after controlling for the effect of litter size on gestational weight gain, which also influenced maternal weight gain throughout pregnancy (beta=4.8920, t(94)=11.978, *p*<2.2e-16). There were no significant differences between females mated with PS-F0 and PC-F0 males in terms of frequency of maternal licking (t(182)=−0.543, *p*=0.588; **Figure 2c**). However, females that mated with stressed males showed reduced frequency of nursing during the first postnatal week (Beta=−0.06, t(183)=−2.227, *p=*0.0272; **Figure 2d**). These effects were driven by a reduction of nursing during postnatal days 4-5 (beta=−0.129, t(34)=−2.445, *p=0.02* and beta=−0.10, t(34)=−2.011, *p=*0.05, respectively).

**Figure 2.**
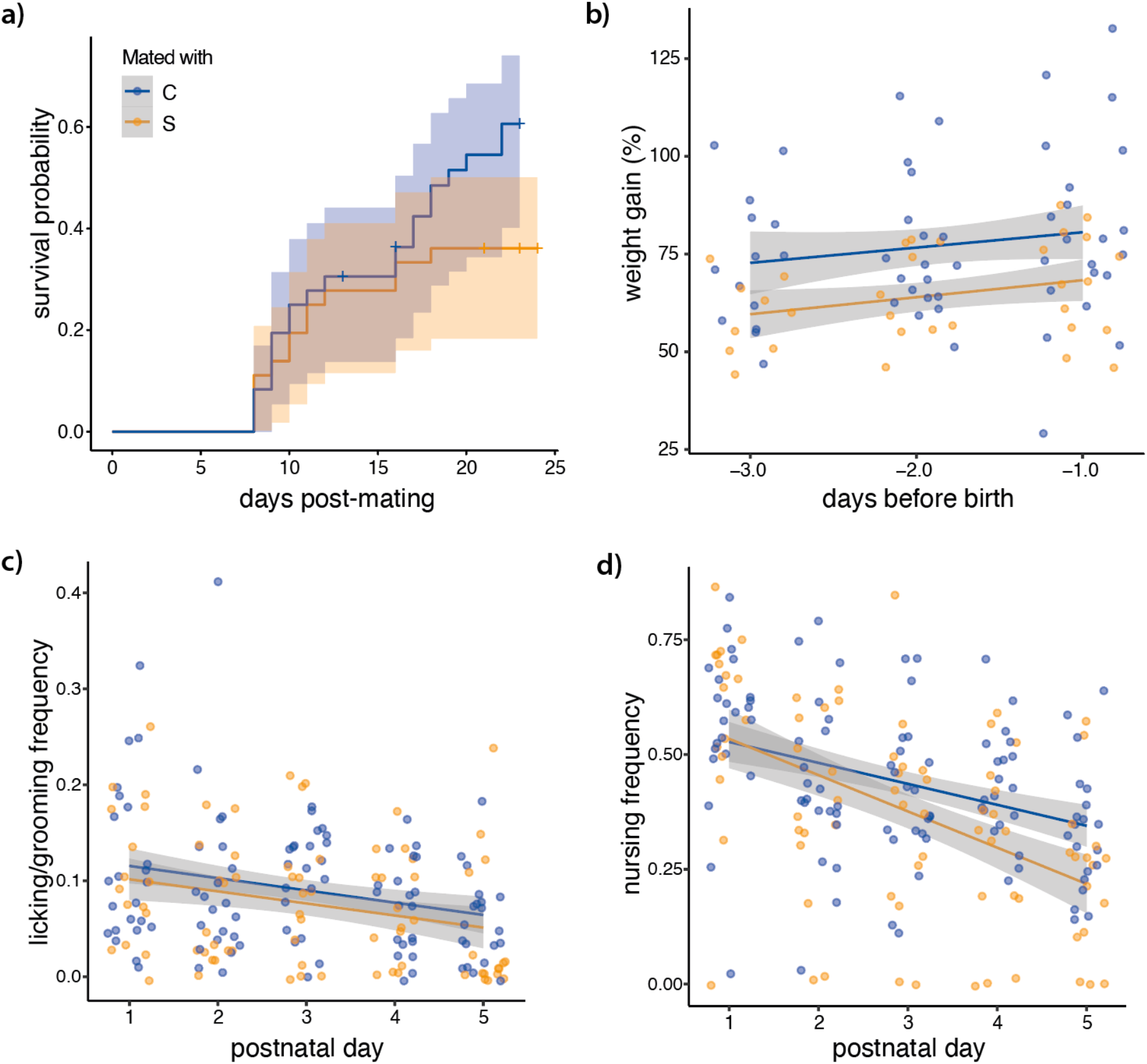
Females mated with stressed (S; green) males have **(a)** less successful pregnancies. **(b)** gain less weight during gestation and **(c)** nurse offspring at reduced frequencies compared to females mated with control (C; blue) males. Mate condition did not affect female licking and grooming of pups.

### Offspring behavior

#### Open-Field Test

The effects of paternal stress on offspring open-field behavior were sex-specific. When males and females were analyzed separately we find that paternal stress resulted in male offspring that spent more time (beta=50.42, t(29)=2.141, *p=*0.041; **Figure 3a**) and travelled a greater distance (beta=2.1642, t(29)=2.261, *p=*0.031) in the center of the open-field arena. There were no effects of paternal stress on female offspring in the open-field test.

**Figure 3.**
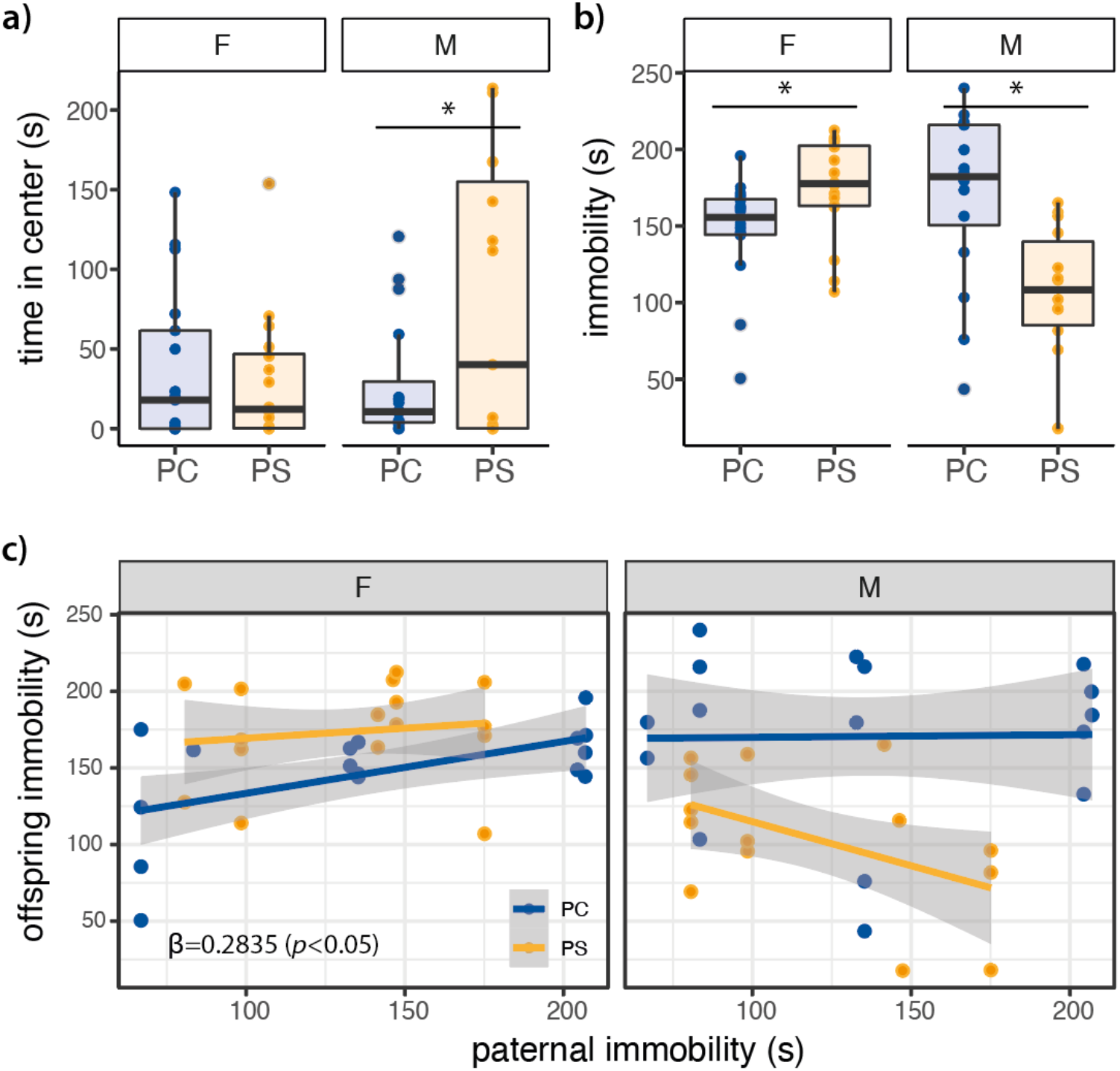
Paternal stress (PS) results in sex-specific behavioral outcomes in offspring compared to control fathers (PC). **(a)** Male PS offspring show increased time spent in the center of an open-field. **(b)** Female PS offspring show increased mobility in a forced swim test, whereas male immobility is reduced. **(c)** Fathers’ immobility is positively correlated with immobility score of daughters but not sons in the forced-swim test (*\*p<0.05*).

#### Forced-Swim Test

The effects of paternal stress on offspring force-swim behavior were also sex-specific. Female offspring of stressed males spent less time swimming passively (beta=−26.52, t(30)=−2.183, *p*=0.037) whereas males spent more time swimming passively (beta=58.67, t(30)=−3.291, *p*=0.0027). Male offspring of paternally stressed males also spent more time actively swimming/struggling (beta=3.8288, t(28)=2.671, *p=*0.013) whereas no such difference were found within female offspring (beta=−0.4875, t(30)=−1.297, *p*=0.20). Moreover, female offspring of paternally-stressed males show increased immobility (beta=28.67, t(28)=2.523, *p*=0.02), whereas male offspring immobility was reduced (beta=−67.67, t(28)=−3.298, *p=*0.01) during the forced-swim test (**Figure 3b**). Interestingly, there was a positive correlation between the duration of time spent immobile by father’s and daughters, which in addition to the paternal stress condition, independently and positively influenced female offspring (beta=0.2835, t(28)=2.330, *p*=0.03). There was no such influence of fathers’ immobility on immobility in male offspring (**Figure 3c**).

### Offspring gene expression

Paternal condition had significant interactive effects on brain-derived neurotrophic factor (*Bdnf*) and corticotropin releasing hormone (*Crh*) expression in the developing hypothalamus. Hypothalamic *Crh* mRNA levels on postnatal day 6 were increased in female (beta=0.3282, t(14)=2.257, *p*=0.04) but not male (beta=0.21, t(14)=1.585, *p*=0.135) offspring sired by PS fathers (**Figure 4a**). Hypothalamic *Bdnf* levels, however, were significantly reduced in male (beta=−0.23, t(14)=−2.056, *p=*0.05) but not female (beta=0.11, t(14)=1.443, *p=*0.17; **Figure 4b**) PS offspring at the same time point. These effects did not persist into adulthood as there were no differences in either *Crh* or *Bdnf* expression in the adult hypothalamus of offspring of either stressed or control fathers of either sex, nor was there any interaction [full model for *Crh*: (F(3,28)=1.599, p=0.92), full model for *Bdnf* : F(3,28)=1.051, p=0.385)].

**Figure 4.**
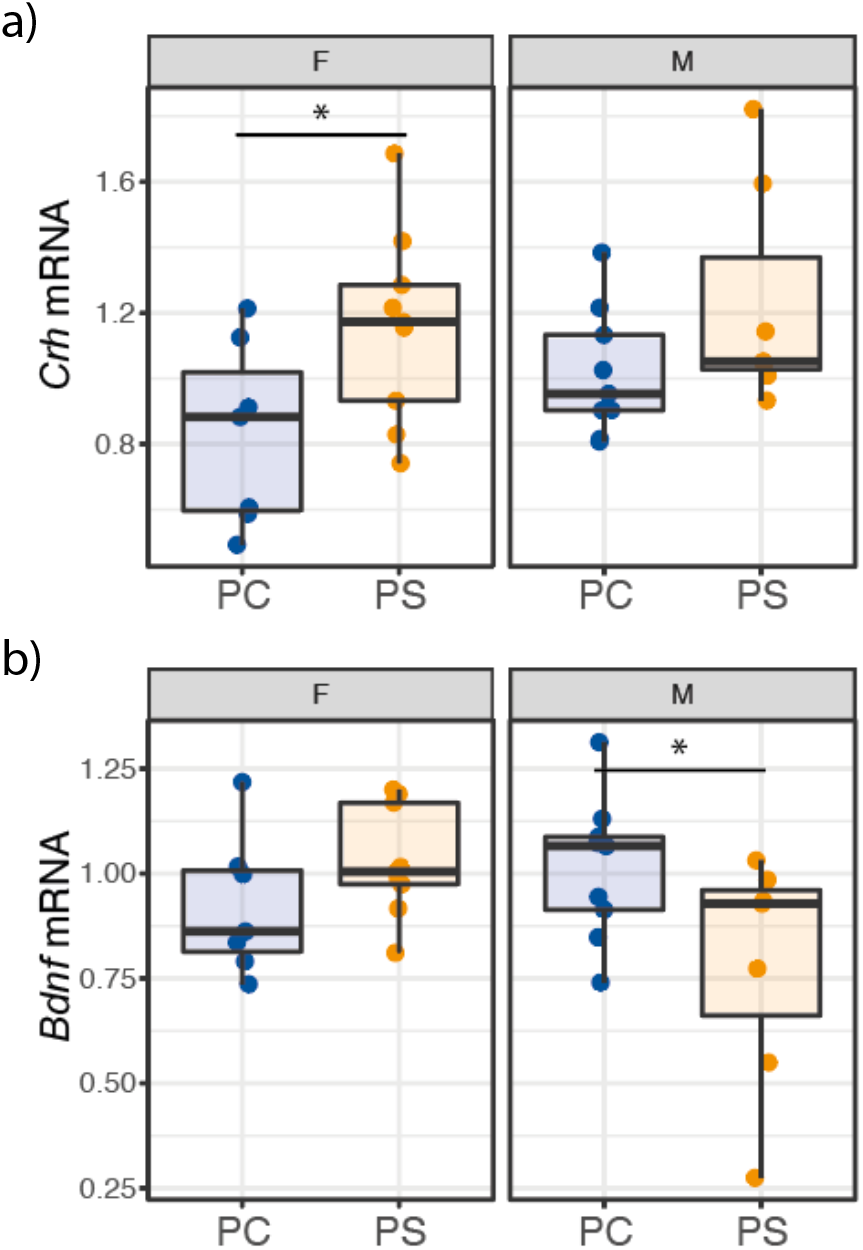
Gene expression of **(a)** corticotropin-releasing hormone (CRH) is increased in the developing (PN6) hypothalamus of female offspring of stressed fathers. **(b)** mRNA levels of BDNF are decreased in male offspring. (\**p*<0.05)

### Mediation of paternal effects by mothers

Given that paternal stress altered both maternal behaviors of mates as well as offspring outcomes, we tested if paternal effects were mediated, at least in part, by changes in maternal behavior. As described above, paternal stress condition was a significant predictor of offspring immobility in the FST in both sexes (Total effect). Both prenatal weight gain (beta = 21.19 (−.48 – 57.52), *p*=0.05) and postnatal nursing (beta= 19.96 (−4.26 – 44.98), *p*=0.05) were significant partial mediators of this effect in male offspring (Indirect effect; **Figure 5**). No such mediating relationship was found in female offspring (**Figure 5;** see **Table S2**).

**Figure 5.**
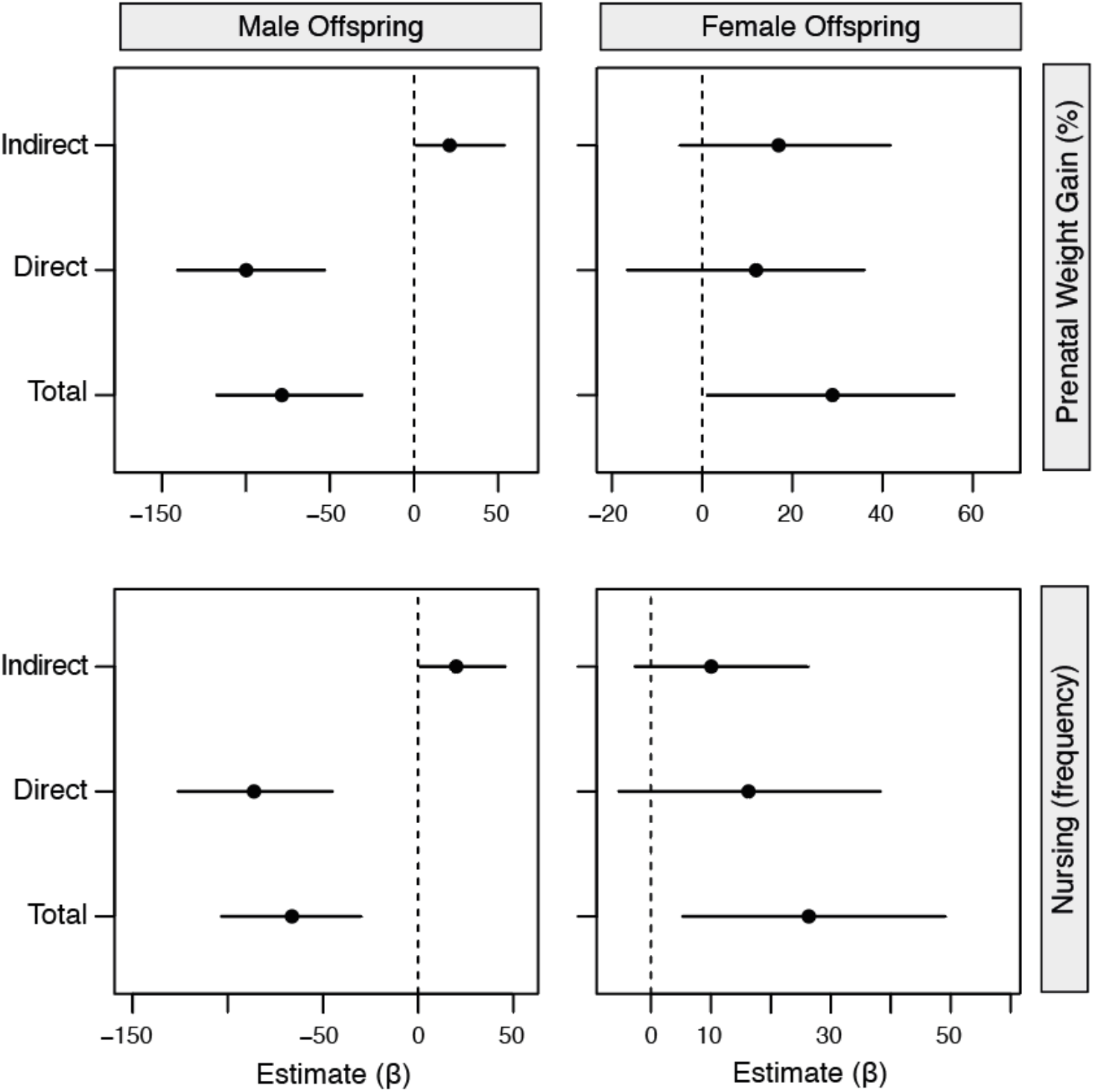
Forrest plot of slope estimates (β) for mediation analyses predicting the effect of paternal stress condition on offspring duration of immobility in the forced swim test (lines represent 95% Confidence Intervals). There was a significant mediating effect of maternal investment (prenatal weight gain and nursing; Indirect effect) in male offspring behavior. This was in addition to a significant direct effect of fathers on offspring and total effect (including maternal variables) indicating a partial mediation in males. No such mediating effect was found in females.

## Discussion

The current study demonstrates that the effects of paternal stress on offspring development are sex-specific. Males that are chronically stressed sire female offspring that show increased anxiety and increased depression-like behaviors. This paternal effect was associated with increased expression of *Crh* mRNA in the developing hypothalamus. Paradoxically, male offspring of stressed fathers show reduced anxiety-like and depression-like behaviors as adults. Moreover, we show that the effects of paternal stress are partially mediated via mothers’ changes in maternal investment in response to their mates. Taken together, these data suggest a role for maternally-induced effects in propagating the effects of paternal stress in addition to any direct effects paternal condition may have on offspring development.

### Sex-specific effects of paternal stress on offspring development

Our findings indicate that paternal stress in Balb/C mice results in an increase in anxiety- and depression-like behavior in female offspring. However, males show a pronounced reduction in these behaviors compared to offspring sired by control males. Overall, these results suggest that male offspring of stressed fathers are less sensitive to stressors (induced by open-field and forced-swim) and more active/exploratory. Consistent with this suggestion, we see increased *Crh* and decreased *Bdnf*. Given that *Crh* is involved with increased stress-reactivity of the hypothalamic pituitary adrenal axis (30,31) and *Bdnf* maintains energy homeostasis and reduces stress reactivity (9,32), these data are consistent with divergent stress programming in response to paternal stress between the males and females. This interpretation is consistent with previous work in three-spined sticklebacks suggesting that paternal stress may prime sons for riskier environments. For example, in three-spined sticklebacks, paternal predation exposure resulted in sons that were more active and exploratory, which resulted in more risk taking and reduced survival when confronted with a predator (33).

While there have been many studies showing sex-specific effects of paternal stress on offspring development, there has been no consistent indication regarding the direction and magnitude of effects on offspring phenotype (7,8,11,34). For example, males stressed early in life (maternal stress combined with maternal separation) sired offspring (both male and female) that exhibited reduced anxiety-like behavior across a battery of tests with no effect on depression-like behavior (8,14). However, chronic variable physical stress during adolescence or adulthood had no effect on baseline anxiety or depression-related behaviors (11). In contrast, social defeat stress in adulthood, resulted in elevated levels of anxiety- and depression-like behavior in both males and females, with more pronounced effects in males (7,35). These seemingly paradoxical findings are likely to be due to a combination of stressor timing (*i.e.*, the developmental stage when stress was experienced), the qualitative nature of stressor and the duration of exposure (i.e., short vs. long-term exposure). For example, different stressors at different time points may have varied effects on sperm development depending on the stage of spermatogenic cycle affected, which could, in turn, affect sperm content and quality at fertilization (36). Another possibility is that non-genetic paternal factors may interact differently depending on the genetic backgrounds of mice. Much of the work described has been in C57Bl6 mice, whereas here we use Balb/C mice which are generally less social and more sensitive to stress (37,38).

Despite these consistent reports of sex-specific effects on offspring development, we still do not have a clear understanding of how these effects arise mechanistically. Suggested explanations in the literature include sex-chromosome linked paternal epigenetic variation (39,40). Moreover, given differences in sex hormone release and sex-specific epigenetic programming events in utero (41,42), there may be differences in timing that render one sex more or less sensitive to paternal-associated variation. Relatedly, there may be differences in the provision of postnatal maternal care, which could further induce sex-specificity in behavioral outcomes (43,44). Given that females are more vulnerable to stress-related disease (45), this is a key area for future work.

### Paternal effects via the germline

Paternal effects on offspring development are particularly intriguing because they highlight the opportunity for environmentally-acquired epigenetic marks and signals to be inherited across generations. Though DNA methylation was initially identified as a potential candidate, it is increasingly considered as a less robust heritable non-genetic mark (5,14). This is primarily attributed to the major waves of reprogramming during development that erase any acquired DNA methylation and render transmission across the germline rare (41). More recent work on paternal stress has focused on sperm RNAs which could be transferred at fertilization from the sperm nuclei itself, or hitchhike via extracellular vesicles that are fused to sperm (8,12,13,35,46,47). Critically, we and others have shown that artificial reproduction (e.g., embryo transfer and in vitro fertilization) are sufficient for the transmission of paternal experience on offspring (4,7,8,35). For example, stressful experiences of males (both in early-life as well as adult exposure) result in changes in small and long noncoding RNAs in sperm, which when transmitted *via* in vitro fertilization (IVF) influences offspring phenotype in a sex-specific manner (7,8). These data are suggestive of a causal role with evidence that these RNAs may bind to consensus sequences in the developing embryo to influence transcriptional programs (6).

### Role for paternally-induced maternal effects

Though the mechanistic basis of paternal effects has solely focused on the transmission of non-genetic marks and signals *via* sperm, we have previously shown that paternally-induced maternal effects might indirectly mediate, at least in part, some phenotypic transmission. This study further adds to that concept, showing that mating with stressed males leads to a reduction in maternal investment (both pre- and postnatal) in Balb/C mice with repercussions for offspring developmental trajectories. Our results indicate that although paternally stressed fathers had direct effects on offspring behavior, the strength of these effects were partially mediated through mothers’ change in behavior. Previously, we showed that mating with socially-enriched Balb/C mice or food-restricted C57Bl6 male mice resulted in increased maternal investment (4,23). Using embryo transfer, we showed that while food-restricted fathers could directly influence growth rate, hypothalamic gene expression and behavior in female offspring, many of these phenotypes are absent or reversed under natural mating conditions. We further showed that this was likely due to increased maternal investment in response to food-restricted mates, which occurs only when females mate naturally with food-restricted males (rather than gestate transplanted embryos) (4). The finding that the effects of chronic social defeat stress in isogenic male mice are not completely transmitted to offspring when sired using IVF lends further support to the possibility that maternal mediation of these effects may play a role (7).

Critically these data suggest that maternal investment can both perpetuate or compensate for male phenotype depending on the nature of the experience and genetic background of adult male mice. It is, therefore, not surprising that different types of stressors, such as chronic physical stress used in this study, may then result in reductions in maternal investment. This effect could result from differences in female assessment of male quality, or sexual interactions at mating or changes in seminal fluid that could prime reproductive hormones (2,20,48,49). Regardless of how these effects emerge, these data add to a growing body of work suggesting that these paternally-induced maternal effects can additionally shape the direction and magnitude of phenotypic change in response to paternal phenotype.

### Conclusions

The idea that environmentally-induced signals could be inherited *via* the germline has provoked re-evaluation of our definitions of heritability. In the current study, we show how social interactions between parents may provide an additional route through which paternal experience may influence offspring development, even when fathers’ do not provide parental care themselves. While this has been well-documented in non-mammalian species, we have shown this to occur in response to paternal social isolation/enrichment (23), dietary restriction (4) and now paternal physical stress in inbred laboratory mice indicating that this is a robust phenomenon with implications for offspring developmental trajectories. Therefore, there are multiple pathways through which the experiences and life-histories of parents interact to drive phenotypic variation which can impact the subsequent direction and strength of transmission of parental effects. Predictions about the long-term heritability of epigenetic effects should take these additional sources of variation into account.

## Supplementary Tables

**Table S1.**
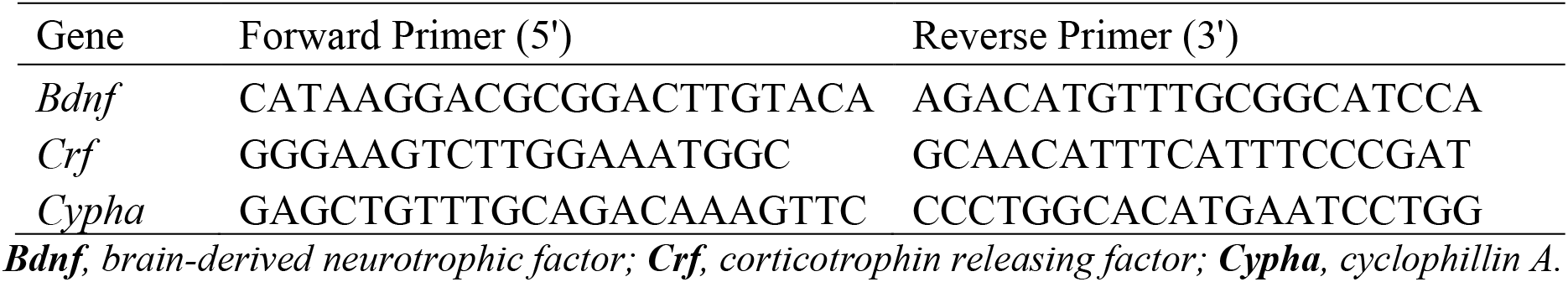
Primer sequences for qPCR

**Table S2.** Mediation analysis of maternal weight gain and maternal nursing on offspring immobility in the forced-swim test.

|  | Estimate | 95% CI Lower | 95% CI Upper | *p* |
| --- | --- | --- | --- | --- |
| *Male Offspring - Maternal Weight Gain* |  |  |  |  |
| ACME | 21.19 | -0.48 | 57.52 | 0.05 |
| ADE | -99.80 | -142.38 | -59.47 | 0.00 |
| Total Effect | -78.62 | -123.74 | -35.57 | 0.00 |
| Prop. Mediated | -0.27 | -1.27 | 0.00 | 0.05 |
| *Male Offspring - Maternal Nursing* |  |  |  |  |
| ACME | 19.96 | 0.09 | 43.23 | 0.05 |
| ADE | -86.26 | -126.62 | -45.75 | 0.00 |
| Total Effect | -66.30 | -98.46 | 29.98 | 0.00 |
| Prop. Mediated | -0.30 | -0.94 | 0.00 | 0.05 |
| *Female Offspring - Maternal Weight Gain* |  |  |  |  |
| ACME | 16.95 | -4.26 | 44.98 | 0.16 |
| ADE | 11.95 | -10.48 | 37.97 | 0.33 |
| Total Effect | 28.90 | 3.90 | 59.18 | 0.02 |
| Prop. Mediated | 0.59 | -0.28 | 1.83 | 0.16 |
| *Female Offspring - Maternal Nursing* |  |  |  |  |
| ACME | 10.06 | -3.69 | 25.39 | 0.15 |
| ADE | 16.27 | -8.69 | 37.81 | 0.14 |
| Total Effect | 26.33 | 1.99 | 51.85 | 0.03 |
| Prop. Mediated | 0.38 | -0.38 | 2.06 | 0.16 |

